# State of pedestrian road safety in Uganda: are interventions failing or absent?

**DOI:** 10.1101/736488

**Authors:** Jimmy Osuret, Stella Namatovu, Claire Biribawa, Bonny E Balugaba, Esther Bayiga Zziwa, Kennedy Muni, Albert Ningwa, Frederick Oporia, Milton Mutto, Patrick Kyamanywa, David Guwatudde, Olive Kobusingye

## Abstract

**Introduction:** In Uganda, pedestrians are the most frequently injured category of road users, accounting for 40% of road traffic fatalities and 25% of serious injuries every year. There is paucity of information on existing pedestrian interventions and challenges that affect their implementation in Uganda. In this paper, we ascertain the state of pedestrian road safety interventions in Uganda and explore the challenges in the process of design, implementation, monitoring and evaluation of existing interventions.

**Methods:** We conducted a qualitative study that started with a desk review of existing policy documents, police statistics, media reports, non-governmental organization reports, and published research. We supplemented the review with 14 key informant interviews and 4 focus group discussions. Participants were drawn from various agencies and stakeholders responsible for road safety. In total, we collected and synthesized data on the design, implementation, and evaluation of pedestrian safety interventions from 25 documents. Data were analyzed using qualitative thematic content analysis.

**Results:** The National Road Safety Council within the Ministry of Works and Transport is the lead agency tasked with coordinating all road safety efforts, while the Uganda Police is largely engaged in enforcing pedestrian safety. We identified several existing policies and regulations for pedestrian safety like the Non-Motorized Transport policy whose implementation has been inadequate. Implementation is constrained by weak institutional capacity and limited resources. Moreover, road safety stakeholders operated in silos and this hindered efforts to coordinate pedestrian safety activities. Interventions like road designs were implemented with limited reference to any supporting data and therefore did not cater for pedestrian needs.

**Conclusion:** There are interventions targeting pedestrian safety in Uganda, but effective implementation is lacking or failing due to constraints related to weak institutional capacity. This necessitates strategies to mobilize resources to strengthen the capacity of the lead agency to effectively coordinate road safety interventions.

## Introduction

Road safety receives inadequate attention, yet every year, the burden remains high at 1.35 million deaths and up to 50 million injuries globally(1). The burden of road traffic injuries and deaths is more pronounced among vulnerable road users, especially those living in low-and middle-income countries (LMICs) (1–3). More than half of the global road traffic deaths are among pedestrians, cyclists and motorcyclists who are neglected in road safety management programs in many countries(1). Each year, more than 351,000 pedestrians and cyclists lose their lives on the world’s roads(1). Moreover, between 2013 and 2016, no reductions in road traffic deaths were observed in any low-income country, while some reductions were observed in 48 middle and high income countries(1). Countries that have succeeded in addressing pedestrian road safety have achieved this through implementing a holistic road safety approach that encompasses infrastructure with provision for all categories of road users, a ‘forgiving’ road environment, consistent enforcement of road and vehicle safety standards and regulations, promoting safe road user behaviour, and post-crash care(1).

The road traffic death rate in Uganda is still unacceptably high, estimated at 29 deaths per 100,000 population compared to the global death rate which has remained fairly constant at 18 death per 100,000 population over the past 15 years(1). Pedestrians comprise the largest group of road users killed in Uganda, accounting for about 40% of fatalities and 25% of serious injuries(4). There is pressure for low-income countries like Uganda to address the problem of road traffic crashes with special attention given to all categories of pedestrians and special groups like children, the elderly and persons with disabilities (1). Uganda has a legal framework that underpins pedestrian road safety management under the Non-Motorized Transport policy and the Traffic and Road Safety Act 1998. Road safety interventions elsewhere have been ranked either as proven, promising, or having insufficient evidence in terms of improving pedestrian safety(5). In Uganda, interventions include pedestrian sidewalk, over passes, road safety campaigns and enforcement by Police on speed limits, and road user behaviour(4). However, progress in reducing the incidence of pedestrian road traffic injuries (RTIs) and deaths has been suboptimal, partly because the needs of pedestrians are often not catered for in the planning, design and operation of roads. Other factors associated with pedestrian injuries and deaths include; speed; inadequate pedestrian infrastructure; risk road use behaviour among pedestrian; poor visibility; age (e.g. young and elderly); driving under influence of alcohol; poor road condition; inadequate road safety enforcement; and driver distraction(6, 7).

Considerable research exists on a narrow range of pedestrian interventions and is largely focused on the magnitude, trends and patterns of pedestrian fatalities and injuries (8–12). Stimulating country action to address the problem of pedestrian safety requires an understanding of the wider policy environment and intervention impediments before prioritizing interventions and creating a plan of action. A clear understanding of the policies, guidelines, rules and regulations as well as contextual factors related to politics, environment, economics and capacity is needed for better design of effective pedestrian-targeted interventions in Uganda. This study ascertains the state of pedestrian road safety in Uganda, and explores key challenges in the process of design, implementation, monitoring and evaluation of existing interventions.

## Methods

### Study design

We conducted a qualitative study that utilized an ethnographic approach to understand the policy and programmatic aspects that underpinned existing road safety interventions in Uganda, with focus on pedestrians.

### Sampling and data collection

Data were collected concurrently using three primary methods; document review, key informant interviews (KIIs) and focus group discussions (FGDs). The document review included documents from government, the private sector, government parastatals, and non-governmental organizations (NGOs) and international agencies working on road safety. Documents for review were provided by key informants in hardcopy, and softcopies were downloaded from websites of relevant institutions. Only documents with sections relevant to pedestrian road safety were included in the study (Figure 1 and Table 1). An inventory of all included documents was created for tracking purposes. Two reviewers then independently extracted data from all included documents using structured data extraction forms. The review focused on leadership and stakeholder engagement particularly examining their focus areas, interests, resources and relationships of various stakeholders and their current roles in road safety. In addition, we reviewed existing plans, policies and programs. We extracted data on pedestrian interventions, intervention implementation, and monitoring and evaluation.

**Figure 1.**
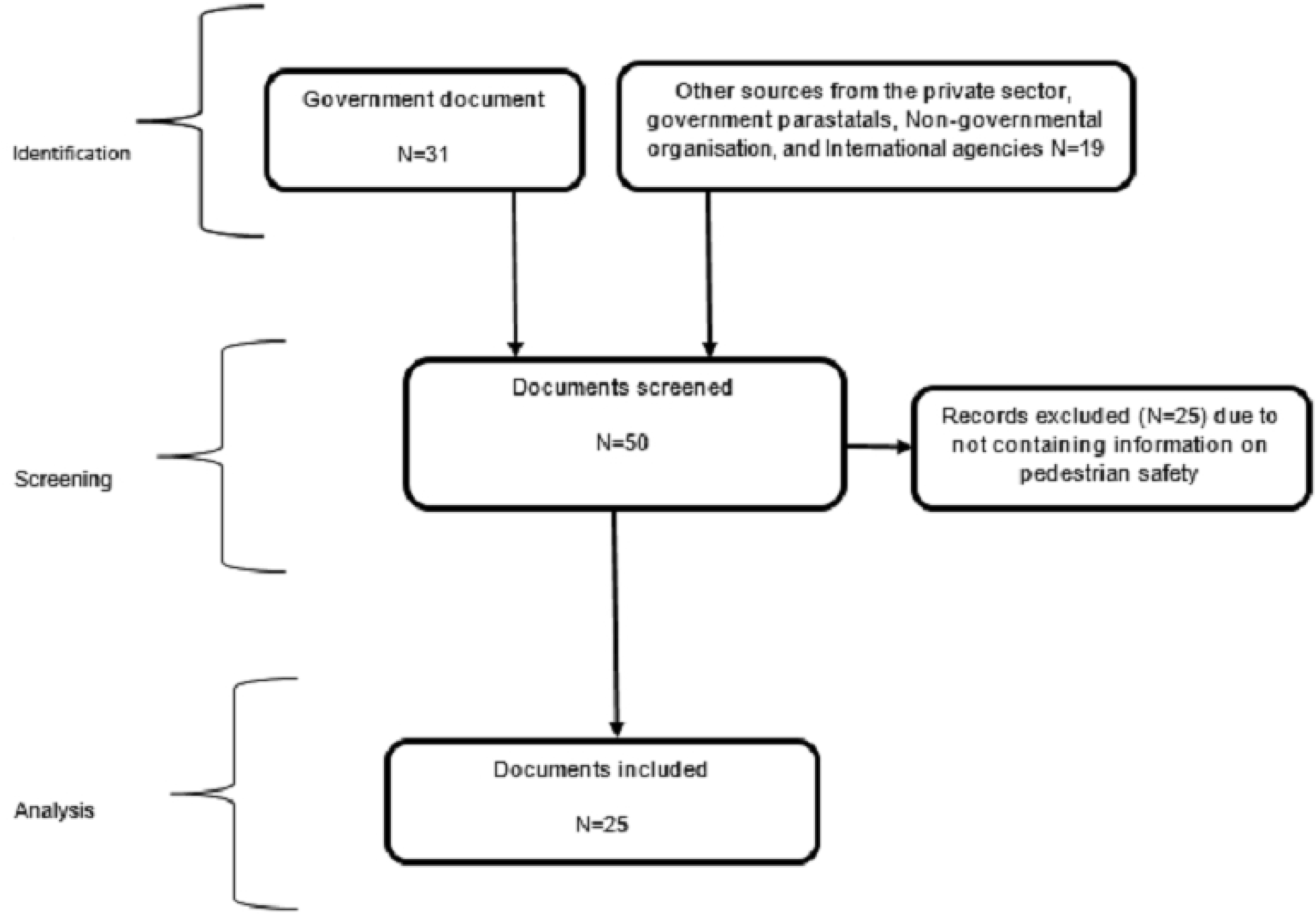
Flow diagram illustrating the document selection process.

**Table 1.** Category of documents included for the review.

| Source | Name of document |
| --- | --- |
| Insurance Regulatory Authority | ✓ Motor vehicle insurance (third party risks) Act 1989 |
| Kampala Capital City Authority (KCCA) | ✓ Kampala physical development plan September 2012<br>✓ The study on Greater Kampala road network and transport improvement in the republic of Uganda November 2010 |
|  | ✓ A detailed strategic implementation plan for the national transport master plan including the greater Kampala metropolitan area 2015-2023<br>✓ National transport master plan including a transport master plan for the greater Kampala metropolitan area (NTMP/GKMA) August 2009 |
| Ministry of Lands, Housing and Urban Development (MoLHUD) | ✓ National physical planning standards and guidelines 2011 |
| Parliament | ✓ Uganda's Road Safety Legislative Action Plan 2018 |
| Uganda Police Force (UPF) | ✓ The Traffic and Road Safety Act, 1998<br>✓ The highway code March 2009 |
| Ministry of Works and Transport (MoWT) | ✓ National road safety policy June 2017<br>✓ International Road Assessment programme Uganda 2010 technical report<br>✓ Ministry of Works and transport, Strategic plan (2011/12—2015/16)<br>✓ Non-motorized transport policy October 2012<br>✓ The traffic and road safety act ,1998 statutory instrument 361-10<br>✓ The traffic and road safety (rules of the road) regulations, 2004.<br>✓ The traffic and road safety (speed limits) regulations, 2004 |
| Uganda National Roads Authority (UNRA) | ✓ The Uganda national roads authority (general) regulations, 2017.<br>✓ General specifications for road and bridge works |
| Safe Way Right Way | ✓ Road safety quarterly news bulletin for Safe Way Right Way Uganda, issue no. 006: February 2016<br>✓ Road safety quarterly news bulletin for Safe Way Right Way Uganda issue no. 005: October 2015<br>✓ Safe Way Right Way quarterly newsletter: Jul-Sept 2017<br>✓ Road safety inspection Kampala-Malaba highway (Document not dated)<br>✓ Concept note: induction of the Parliamentary Forum for Road Safety (PAFROS) – December, 2017.<br>✓ Road safety inspection of Kampala-Hoima road October 2015 |
| United Nations | ✓ Road safety performance review Uganda February 2018 |

In-depth interviews were conducted with 14 purposively selected key informants using an interview guide. The key informants were drawn from stakeholders involved in pedestrian safety, and these included representatives from the Ministry of Works and Transport; Ministry of Lands, Housing, and Urban Development; Ministry of Education; Ministry of Health; the Uganda Traffic Police; the National Road Safety Council; Kampala Capital City Authority, Parliament, and road safety NGOs. We conducted 4 homogenous focus groups with one group each for pedestrians, commuter taxi drivers, boda-boda (commercial motorcycle) drivers, and private car drivers. Focus group participants were purposively selected using convenience sampling. The guides were pretested incorporating feedback prior to data collection. Data was collected by the investigators and trained research assistants. The interview and focus group guides covered aspects on pedestrian safety interventions, impact of interventions, stakeholders involved in pedestrian safety, factors associated with pedestrian injuries and deaths, and challenges impeding implementation. The interviews and discussions were audio recorded after seeking permission from the participants and field notes taken. Probes were applied based on responses of the participants. We conducted key informant interviews and focus group discussions until no new data was attained and saturation was reached.

### Data management and analysis

For the document review, a harmonized summary was created through consensus between the two reviewers and where there were still areas of disagreement, a third reviewer was consulted. The KIIs and FGDs were transcribed verbatim and cleaned. Where discussions were done in Luganda, these were directly translated into English in preparation for analysis. The transcripts were exported to ATLAS.ti Version 7 software tool for coding and analysis qualitative data. For both the KIIs and FGDs, topical codes were created from the guides while others emerged from the data. The codes were then applied by 2 groups in the study team to the transcripts using a qualitative thematic content analysis approach (13, 14) with categories and themes arising from the data.

## Results

The results are presented in 2 thematic topics from the data analysis namely: the state of pedestrian safety in Uganda and challenges in implementing pedestrian safety interventions in Uganda. The categories and codes from which the themes arose are presented in Table 2.

**Table 2.**
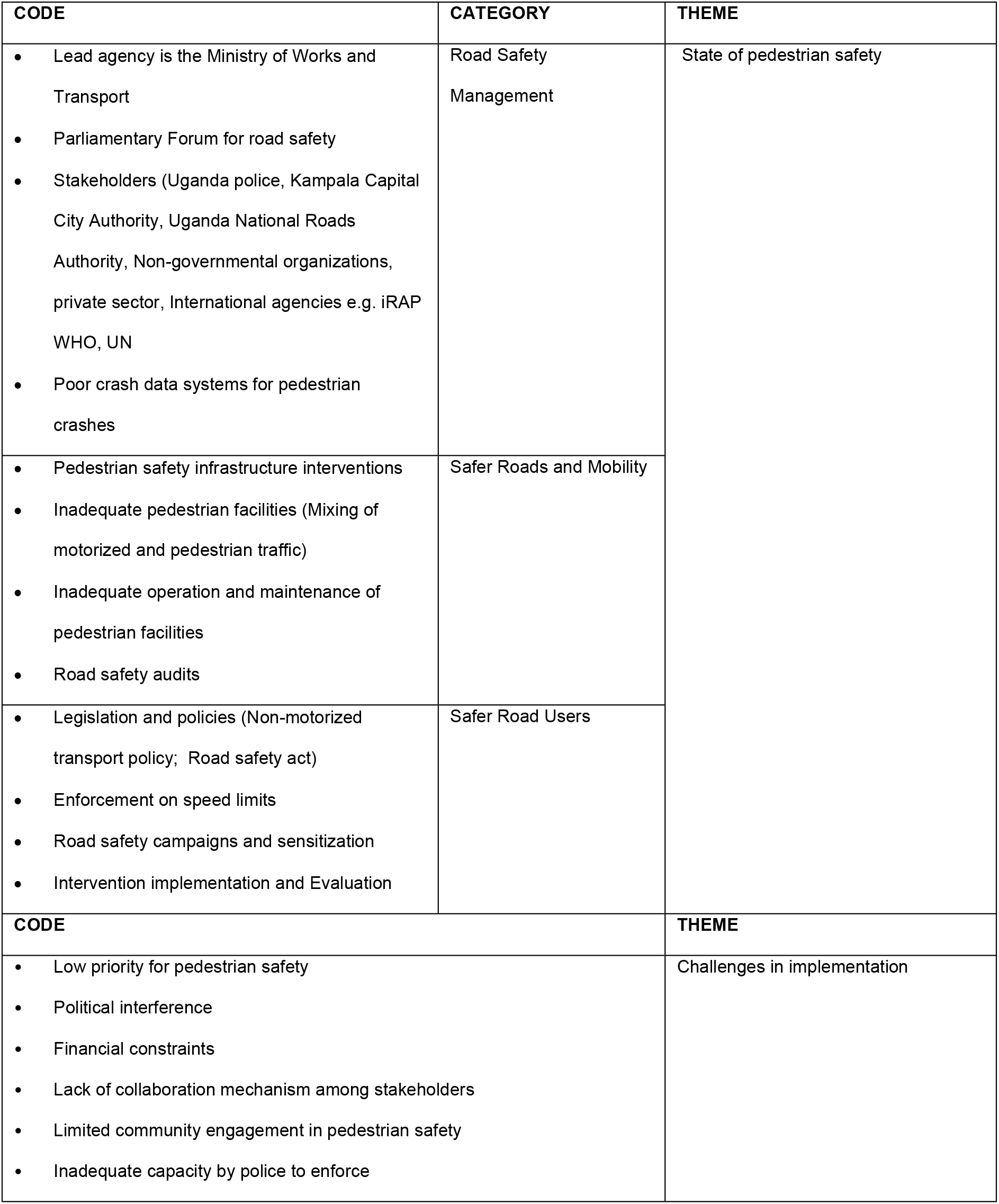
Emerging themes from the Desk review, Key informant interviews and focus group discussions.

### Theme 1: State of pedestrian safety in UGANDA

We identified 3 categories that explained the state of pedestrian road safety in Uganda. Similar codes were categorised under the pillars contained in the global plan for the UN Decade of Action of road safety and they included a) Road safety management, b) Safer roads and mobility and c) safe road users (15, 16)

#### Road safety management

We found that the Ministry of Works and Transport was established as the lead government agency for coordination of all road safety activities operationalized by the National Road Safety Council(17). Uganda also has a parliamentary forum on road safety whose core mandate is to develop legislative action plans on road safety and participation in road safety campaigns. The Uganda Police was reported to be engaged in several enforcement activities such as vehicle inspection and enforcement on the road. We found several stakeholders including the Kampala Capital City Authority, Ministry of Health, Uganda National Roads Authority, civil society, the private sector, NGOs, international organizations who were directly or indirectly involved in pedestrian safety. We noted siloed implementation among the stakeholders with efforts to create multi-sectoral partnerships mostly visible during national events such as the United Nations and the Uganda national road safety weeks. We found that pedestrian safety activities within Kampala were largely done by Kampala Capital City Authority and the Uganda Police playing the key role of enforcement.

> *“…Uganda Police give strategic directives to ensure that it achieves its mandate of reducing crashes and we do it by enforcing regulations educating road users and coordinating with other stake holders to ensure crushes are addressed in the country”*. Key informant
>
> *“It’s the traffic police officer who helps the pedestrians to cross the road”*. Commuter taxi driver—FGD participant

The document review revealed that data systems to support on-going monitoring and evaluation of pedestrian safety do not provide a true estimate of the burden of road traffic crashes, injuries, deaths, and their economic impact. Existing data management systems by the Ministry of Health and the police report different estimates for pedestrian injuries(17)

#### Safer roads and mobility

The pedestrian safety interventions and activities identified from the documents and interviews included operation and maintenance of road infrastructure, road audits, and provision of pedestrian facilities especially within urban areas. However, some roads were poorly maintained, lacked pedestrian crossings and markings, and delays were reported in carrying out periodic maintenance works. In some areas, roads were reported to be narrow with inadequate safe walking facilities. Roads were designed and constructed without considering the needs of pedestrians and other non-motorized modes of transport.

> *“There are planners who think that roads are for vehicles and there are some designers who design with the thinking that roads are for vehicles only”*. Key informant

The pedestrian facilities are also encroached on by other activities like street vending, parking and motorists who drive on the few available pedestrian walkways. Competition for the limited space puts pedestrians at risk.

> *“… Our roads are narrow and congested. For instance, there is mixing of hawkers, boda-boda riders, someone is crossing and as you try to dodge a pothole you knock pedestrians”*. Commuter taxi driver—FGD participant.

#### Safer road users

The Uganda National Road Safety policy and Non-Motorised Transport policy outline priority areas for action to improve road safety for vulnerable road users like pedestrians. The guidelines have provisions for safe pedestrian infrastructure. Pedestrian interventions include provision for pedestrian access routes, prohibition of parking on kerbs, and keeping walkways safe, clear, and well lit. For roads without provisions for pedestrians, it is stipulated that pedestrians walk as far as practically possible from vehicular traffic and against traffic flow. However, implementation of the Non-Motorised Transport policy has been limited. There is also a policy on compulsory insurance against third party risk which is used to make claims for post-crash care for pedestrian victims.

The Uganda Police was reported to be engaged in several enforcement activities like vehicle inspection, blood alcohol content limit and speeding checks. The “Fika Salama” operation (a road safety campaign launched by the Uganda Police in August 2016) was reported to have improved road use discipline although no evaluation on effectiveness was available.

> *“….if the drivers know that the police officer is there, they reduce the speed”*. Commuter taxi driver—FGD participant.
>
> *“When we started Fika Salama [a road safety intervention] you no longer hear people say that the accidents happen on some roads because they are slippery. They now agree that some of the crashes were due to poor road user behaviour”*. Key informant

Sensitization and road safety campaigns were carried out among car drivers, motorcycle drivers, and school children. Children were targeted through their curriculum on road safety because they were willing to learn and were an avenue for passing on pedestrian road safety information to their peers and parents. However, road safety awareness is sporadic and carried out whenever there is a pedestrian crash tragedy or during the national road safety week. The 2017 national road safety week was themed “Think! We are all pedestrians”

> *“… These children of primary school when they learn to respect the road they grow with it [the road safety discipline]”*. Key informant

Uganda has several existing guidelines, rules and regulations (table 1) with a bearing on pedestrian safety that guide implementers during the design of road safety interventions as noted by the key informant. Sources of data that led to the formulation of various interventions include the Uganda Police traffic crash report and statistics from the United Nations and the World Health Organization. However, there are instances where interventions were implemented due to public demand e.g. if a spot has many pedestrian crash incidents then a hump is placed.

> *“…usually when we are planning we use the physical planning standards and these standards have the size of the road, you know that the road should be of a minimum size, And we know that this road is in position to cater for a carriage way, to cater for services and infrastructure and even to cater for the pedestrians walk ways and so on depending on the planning which is available”*. Key Informant
>
> *“If for example there is an accident spot and people are complaining about it many times, we come in with something [intervention] like a road hump to slow down traffic”*. Key informant

There is no formal monitoring and evaluation mechanism for the effectiveness of existing pedestrian safety interventions.

> *“…there is quite some work to do in that area, we don’t have very robust monitoring and evaluation. All we know is that when we do some intervention we get some feedback from the public that now the danger has been averted”*. Key informant

### Theme 2: Challenges in implementation of pedestrian road safety interventions

Pedestrian safety is of low priority considering other public health threats and therefore vulnerable road users receive inadequate consideration during planning and resource allocation for interventions.

> *“Government priority for road safety is still low. Let me tell you, about 30 or more people died last week in crashes. If these were from nodding disease [a disease that has affected children in parts of northern Uganda], Parliament would be up in arms for money for nodding disease”*. Key informant

Political interference was also identified as a deterrent in enforcement and implementation of pedestrian interventions. One of the participants reported that there were instances where

> *“…there are scenarios where the enforcement officers can go [to enforce road safety regulations], and they are not allowed to do that [by the politicians]”* Key informant

The document review revealed weak institutional framework and low capacity at almost every level and this hindered implementation of many policies and regulations like the Non-Motorised Transport policy. Implementation for some pedestrian safety interventions was reported to have been done partially. Limited financial resources allocation was the major hindrance to the implementation of pedestrian safety interventions and policies.

> *“The most common one [hindrance] would be finances because with road safety you need a lot of finances - you need posters, you need fliers, you need to write the message…”*. Key informant

The National Road Safety Council has limited capacity to coordinate all road safety activities including provisions for vulnerable road users. The lead agency did not have a concrete multi-sectoral action plan, and there were no targets for the reduction of pedestrian injuries and deaths in the country. In some instances, various stakeholders involved in pedestrian safety were reported to duplicate interventions already being implemented by others.

> *“The challenge we get is that some of the interventions are not coordinated (hmmm) so you have this one [stakeholder] is doing something similar to another, so the programs are not coordinated. They all compete for visibility”*. Key informant

Community involvement in decision making about pedestrian road safety interventions was minimal as reported from the focus group discussion. Some interventions were implemented without community involvement and consultation and this affects their adoption.

> *“There is a flyover which was put in Nakawa road for pedestrians to use but since they were not sensitized about its importance, they don’t use it; they all use the road. The same applies to the Kalerwe roundabout, the pedestrians use the road yet a flyover is there, but generally, it was not well positioned, it would have been [better] near the market”*. Commuter taxi driver—FGD participant.

Document reviews indicated inadequate capacity and lack of equipment for the National Road Safety Council, other government agencies, and the police to implement and enforce pedestrian safety

## Discussion

We found existing guidelines, regulations, and policies on pedestrian safety and these were reported to inform intervention design and implementation of road systems to a small extent. There were instances where public outcry on pedestrian crashes prompted implementation of traffic calming measures like humps. The existing sources of pedestrian crash information for designing interventions do not provide a true estimate of the burden of road traffic crashes, injuries, deaths, and their economic impact [10]. This is because the road safety data sources from police, hospitals and mortuary cannot be linked together. In LMICs like Uganda, road traffic data is collected from multiple sources which suffer data quality issues of completeness and underreporting(18). Data quality concerns are common in LMICs and yet the number of pedestrian RTIs and deaths is high in these countries (1, 19). This is not the case with high income countries that have invested in credible data systems(1).

The lack of a formal mechanism to guide in the design, implementation, monitoring and evaluation of intervention effectiveness is an indication that roads are designed and constructed with limited consideration given to the needs of pedestrians and other non-motorized modes of transport. The rationale in the design and implementation of road systems in high income countries is to provide safety of all road users and traffic management(20). The finding has implications for establishment of a formal mechanism to guide intervention design and implementation. There is need to monitor and evaluate intervention effectiveness using credible data sources as well as to document and disseminate good practices in the reduction of pedestrian injuries(15, 20).

There is evidence of reduction in pedestrian crashes especially when pedestrians are separated from motorised traffic(21). However, existing engineering measures in Uganda like pedestrian sidewalks, and overpasses have shown limited evidence on improving pedestrian safety. Many times, the needs of pedestrians are not catered for in the planning, design and operation of roads in many LMICs (21). Pedestrian facilities in these countries are inadequate and poorly maintained. These are consistent with our findings, where some roads were reported to be narrow and lacking safe walking facilities to accommodate pedestrian volumes. As a result, pedestrians are forced to walk in the same space as motorised traffic which increases their risk of injury and death. This calls for availing of sidewalks and crossing points in areas where pedestrian volume and risk is highest.

The major barrier to implementation of pedestrian safety interventions and policies in Uganda are linked to the challenges of the National Road Safety Council, which is mandated to coordinate all road safety activities in the country. The limited capacity by the lead agency leads to failure in coordination among various stakeholders and hinders community involvement (22). Due to the inadequate coordination among stakeholders, there were instances where implementation of pedestrian safety activities was in silos or duplicated. Findings from other studies show that the absence of a clear empowered lead agency for road safety affects resource allocation which in the long run hinders the implementation of pedestrian safety interventions and policies(1). There is need to strengthen coordination of road safety through multisectoral collaboration and advocating for additional resources for road safety.

We noted that Pedestrian safety was a low priority in light of other public health challenges and often suffered political interference during implementation. This is contrary to high income countries that have political commitment and dedicated institutional effort to manage road safety (23). Dealing with pedestrian RTIs can be achieved through concerted efforts at national level(15). Achieving national pedestrian safety management requires political and economic commitment that is demonstrated through effective institutional leadership within responsible agencies for road safety(1, 23). The challenge in Uganda remains to generate sustained political will and long term investment programs for road safety.

One of the limitations of the study is that we did not use a comprehensive search strategy to identify the documents included in the review and might have missed out some literature on pedestrian safety in Uganda. However, we addressed this by supplementing the desk review with a qualitative component to obtain thick descriptions, utilising key informant interviews and focus groups with people knowledgeable and involved in pedestrian safety.

## Conclusion

The study found existing interventions including a Non-Motorised Transport policy, and guidelines and regulations aimed at reducing the incidence of pedestrian road traffic injuries in Uganda. The lead agency did not have a concrete multi-sectoral action plan, and there were no targets for the reduction of pedestrian injuries and deaths in the country. Many interventions were being implemented with no evidence for their effectiveness, and no formal evaluations. This necessitates strategies to mobilize resources to strengthen the capacity of the lead agency to coordinate the planning and implementation of evidence-based interventions by all key stakeholders involved in pedestrian road safety.

## Acknowledgements

We express special thanks of gratitude to the study participants and research assistants for taking part in the study.

